# Multilevel selection favors fragmentation modes that maintain cooperative interactions in multispecies communities

**DOI:** 10.1101/2021.03.18.436065

**Authors:** Gil J. B. Henriques, Simon van Vliet, Michael Doebeli

## Abstract

Reproduction is one of the requirements for evolution and a defining feature of life. Yet, across the tree of life, organisms reproduce in many different ways. Groups of cells (e.g., multicellular organisms, colonial microbes, or multispecies biofilms) divide by releasing propagules that can be single-celled or multicellular. What conditions determine the number and size of reproductive propagules? In multicellular organisms, existing theory suggests that single-cell propagules prevent the accumulation of deleterious mutations (e.g., cheaters). However, groups of cells, such as biofilms, sometimes contain multiple metabolically interdependent species. This creates a reproductive dilemma: small daughter groups, which prevent the accumulation of cheaters, are also unlikely to contain the species diversity that is required for ecological success. Here, we developed an individual-based, multilevel selection model to investigate how such multi-species groups can resolve this dilemma. By tracking the dynamics of groups of cells that reproduce by fragmenting into smaller groups, we identified fragmentation modes that can maintain cooperative interactions. We systematically varied the fragmentation mode and calculated the maximum mutation rate that communities can withstand before being driven to extinction by the accumulation of cheaters. We find that for groups consisting of a single species, the optimal fragmentation mode consists of releasing single-cell propagules. For multi-species groups we find various optimal strategies. With migration between groups, single-cell propagules are favored. Without migration, larger propagules sizes are optimal; in this case, group-size dependent fissioning rates can prevent the accumulation of cheaters. Our work shows that multi-species groups can evolve reproductive strategies that allow them to maintain cooperative interactions.

**Author summary:** In order to reproduce, multicellular organisms and colonial bacteria fragment into offspring groups. Fragmentation modes in nature are very diverse: e.g. some organisms split into two halves, while others release single-celled propagules. These fragmentation modes can have fitness consequences, e.g. small propagules reduce the spread of deleterious mutants. However, the consequences of different fragmentation modes are not yet well understood for groups of cells containing several metabolically interdependent species, such as complex biofilms. We developed a multilevel selection model to investigate the effect of fragmentation mode on the accumulation of deleterious mutants when groups contain multiple species. In such groups, small propagules are not always a viable strategy, because they may harbor low species diversity. We find that alternative mechanisms, such as migration of cells between groups and group-size dependent fissioning rates, can prevent the accumulation of mutants. We also find that multilevel selection can lead to the evolution of fragmentation strategies that allow multi-species groups to thrive in the face of deleterious mutations.

## 1 Introduction

Reproduction is a fundamental feature of life and the *sine qua non* of Darwinian evolution (Godfrey-Smith 2009; Lewontin 1970; Stearns 1992). Despite its centrality in natural selection, there appears to be no unique optimum approach to reproduction in multicellular organisms (Kondrashov 1994; Pichugin et al. 2017; Roze and Michod 2001; Van Gestel and Tarnita 2017). Whenever individual cells abandoned solitary life to form groups—ranging from loose collectives (Claessen et al. 2014) to colonial and multicellular organisms—they came up with a surprisingly diverse menagerie of strategies for the production of reproductive propagues (Godfrey-Smith 2009).

Many multicellular eukaryotes reproduce by undergoing single-cell bottlenecks. For example, sexually reproducing organisms produce unicellular gametes. Single-cell bottlenecks are also common in plants and animals that reproduce asexually, such as the Amazon molly *Poecilia formosa* (Turner et al. 1980), several weevils of the Curculionidae family (Suomalainen 1969), and many angiosperms predominantly in the Asteraceae, Rosaceae, and Poaceae families (Bicknell and Koltunow 2004). An alternative to single-celled propagules is vegetative reproduction in which the offspring develops from a multicellular propagule. This type of reproduction may involve specialized structures, such as conidia (in fungi) or gemmae (in algae, mosses and ferns) (Hughes 1971), or it may happen simply by budding (e.g., in hydra, Galliot 2012) or by fission (as in some flatworms, Åkesson et al. 2002).

This wide variation in modes of fragmentation is not limited to eukaryotes. Some bacterial aggregations, such as the clusters formed by *Staphylococcus aureus*, reproduce by releasing single-celled propagules (Koyama et al. 1977). Others, such as filamentous cyanobacteria, reproduce vegetatively: dividing cells remain physically connected, and fragmentation of these aggregates creates new chains (Herrero et al. 2016). A single parent individual may also divide into two equally-sized multicellular offspring. This occurs, for instance, in the multicellular collectives formed by magnetotactic bacteria (Keim et al. 2004). Alternatively, one parent may simultaneously give rise to many equally-sized offspring. The 16-celled colonial alga *Gonium pectorale* takes this strategy to the extreme, by dispersing into 16 individual cells (Stein 1958).

Why is there such wide variation in the number and size of reproductive propagules? A research program initiated by Kondrashov (1994) attempted to answer this question by considering the evolutionary advantages of unicellular propagules relative to vegetative propagules. When offspring develop from a single-cell propagule, they are genetically homogeneous. If the propagule carried any deleterious mutation, its phenotypic effects will not be masked or compensated by wild-type cells. Therefore, reproductive bottlenecks ensure that natural selection is more efficient at eliminating deleterious mutations (Bergstrom and Pritchard 1998; Grosberg and Strathmann 1998; Kondrashov 1994; Roze and Michod 2001). Normally, in the absence of genetic recombination, deleterious mutations accumulate in the population in a process known as Muller’s ratchet (Muller 1964), becoming abundant and decreasing the population’s mean fitness. This decrease in fitness is called mutation load (Agrawal and Whitlock 2012; Haldane 1937; Muller 1950). In extreme cases, the resulting maladaptation can lead to severe declines in population size, accelerating the accumulation of deleterious mutations by genetic drift. This positive feedback, which may lead to extinction, is termed mutational meltdown (Lynch and Gabriel 1990). By facilitating the purging of deleterious mutations, reproductive bottlenecks slow down the detrimental effects of Muller’s ratchet (Bergstrom and Pritchard 1998) and reduce mutation load (Kondrashov 1994; Roze and Michod 2001).

This family of models also informs our understanding of the evolutionary transition (Szathmáry and Maynard Smith 1995) from unicellular life to multicellularity. That is because mutations that are deleterious for the group may be beneficial for the mutant cell itself. Mutations of this type (which Roze and Michod (2001) call “selfish mutations”) result in “cheater” cells whose fast growth comes at a cost to their group; they include, for example, cancer cells (Evan and Littlewood 1998). Selfish mutations (and cheaters more generally) instantiate a conflict between the direction of selection at the level of the cell and at the level of the group. Reproductive bottlenecks resolve this conflict by reducing genetic variation among cells within offspring, and distributing the variation among progeny groups. This reorganization of genetic variation makes selection at the level of the group more effective than selection at the level of the individual cell, thus furthering the evolutionary transition to multicellularity (Roze and Michod 2001).

While this research program has been fruitful and insightful, it considers only a limited set of propagule production strategies (viz., the production of a propagule of varying size). Recent research by Pichugin, Traulsen, and collaborators (Pichugin et al. 2019; Pichugin and Traulsen 2018; Pichugin et al. 2017) has started to address the wide variety of strategies that exist in nature. Their models exhaustively analyse the fitness consequences of every mathematically possible partition of a homogeneous group. This approach is promising, but it becomes unwieldy to consider more complex scenarios, such as big groups (which have many possible partitions), within-group density dependence, or the effect of mutations, which, as we have seen, can have important consequences.

All of the models described above focus on single-species groups, such as multicellular or colonial organisms. However, multispecies communities also undergo fissioning and fragmentation. Such communities are particularly common in the microbial world: the majority of microorganisms belong for at least part of their life to multispecies groups, such as mixed biofilms that comprise up to thousands of different species (Elias and Banin 2012). Just like single-species collectives, multispecies microbial communities display a wide variety of modes of fragmentation. Host-associated communities can be vertically transmitted between host generations or they can be recruited from the environment (both of which are exemplified in insect microbiomes, Engel and Moran 2013). They can also be formed by a combination of both, which is the case, for example, for the human microbiome (Funkhouser and Bordenstein 2013). Environmental communities can also form by recruitment (e.g., colonization of marine-snow particles, Datta et al. 2016) or from multicellular fragments which detach from mature communities (e.g., some bacterial biofilms, Claessen et al. 2014).

Even though natural biofilms are heterogeneous communities of different microbial species, organisms within them are well adapted to group life. In mixed biofilms, individuals of different species communicate with each other via quorum sensing (e.g., Moons et al. 2006; Riedel et al. 2001) and engage in cooperative interactions with each other (Elias and Banin 2012). Such interactions include cross-feeding, in which byproducts from nutrients that are metabolized by one species then serve as food source for a second species (reviewed in Seth and Taga 2014). Cross-feeding is so prevalent that many microbes have lost the ability to synthesize essential metabolites and became metabolically interdependent (Tyson and Banfield 2005). This type of metabolic specialization allows communities to perform otherwise incompatible tasks, such as photosynthesis and nitrogen fixation (Johnson et al. 2012).

Much like selfish mutations in multicellular organisms, cheaters in mixed biofilms are manifestations of a conflict between levels of selection. Some researchers have taken the view that group selection predominates in mixed biofilms, going so far as to call them “evolutionary individuals” (Day et al. 2011; Doolittle 2013; Ereshefsky and Pedroso 2015). Others warn that this comparison holds only as an analogy, and that the conflict between levels of selection has not been resolved in mixed biofilms (e.g., Clarke 2016; Coyte et al. 2015; Foster and Bell 2012; Nadell et al. 2009). Although the question of whether group- or individual-level selection predominates in mixed biofilms remains controversial, recent modelling work has started to address which conditions need to be met for multilevel selection to operate in these communities (Roughgarden 2020; Van Vliet and Doebeli 2019).

Here, we build on these previous models and further investigate how the conflict between levels of selection can be resolved in both single- and multi-species groups. In particular, we focus on the question of how the mode of fragmentation of the group affects its ability to maintain cooperative interactions in the presence of recurrent mutations to cheating cell types. We developed an individual based, multilevel selection model (Simon 2010; Simon et al. 2013) of groups consisting of cells that interact mutualisticly. Cells in the group stochastically give birth or die, while groups stochastically go extinct or reproduce. Whenever a group reproduces, it follows a predefined “fragmentation mode” that describes how its constitutive cells are arranged over the offspring groups (Fig. 1). Both the size and the number of offspring groups can vary continuously allowing us to fully investigate how the mode of fragmentation affects the ability of the group to resist mutations.

**Figure 1:**
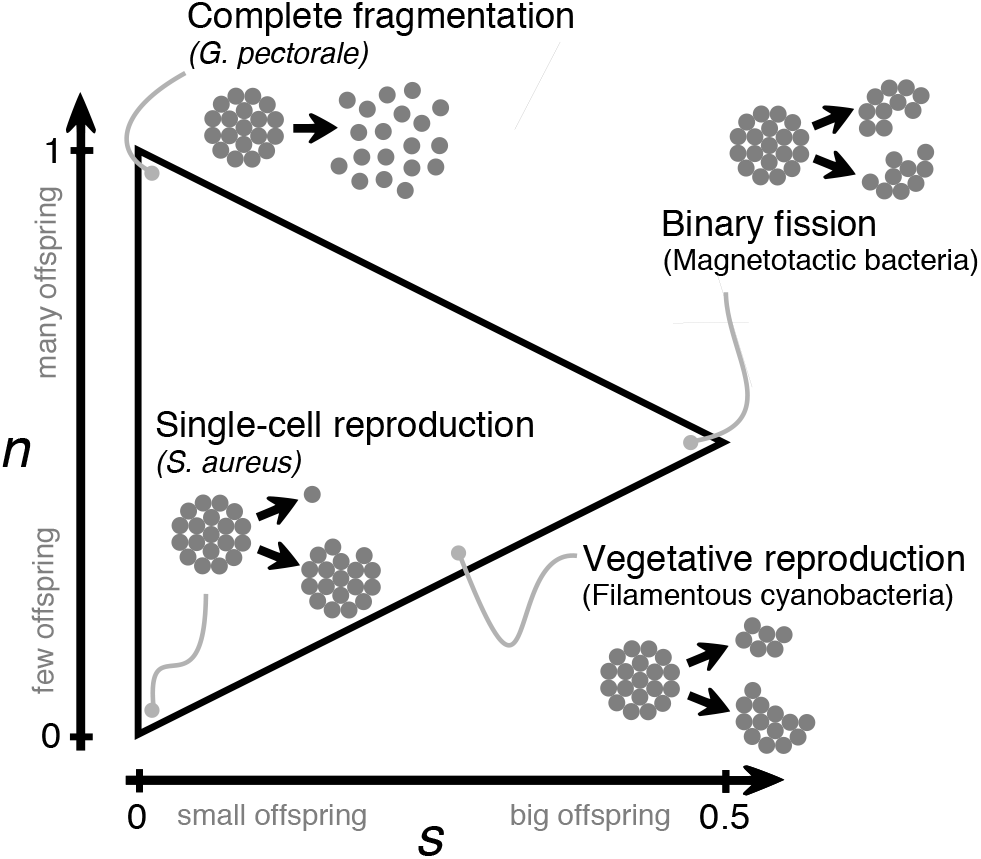
All possible fragmentation strategies in our model can be described using a two-dimensional phase space. Two parameters (*s* and *n*) determine the expected size and number of offspring, respectively (as a fraction of the number of cells in the parent group). Cartoons exemplify some modes of fragmentation. We refer to the three corners of the triangle (complete fragmentation, single-cell reproduction, and binary fission) as archetypal modes of fragmentation.

Drawing from the lessons of models on single-species collectives, we expect that in multi-species groups different fragmentation modes could have strong implications on how well the system can cope with mutants. For example, by decreasing variation within groups, we expect that reproductive bottlenecks intensify group selection and help purge cheater mutants. On the other hand, we expect that the benefits of strong bottlenecks diminish in complex multi-species communities, because small propagules are less likely to contain all of the species that are metabolically interdependent.

Our model confirms these expectations: we find that high-diversity communities face a reproductive trade-off: tight bottlenecks result in the elimination of deleterious mutations, but they also decrease species diversity; the opposite is true when offspring groups are large. Using our model we also investigated strategies that can relieve this trade-off: adding migration between groups reconciles single-cell bottlenecks with the need for diversity, while making group fragmentation rate dependent on group size increases the strength of group level selection thus reducing mutant load. Finally, we extend our model to investigate how group fragmentation rates evolve and we show that groups always evolve to the fragmentation mode that maximizes their resistance to mutants.

## 2 Methods

We developed an individual-based multilevel selection model (Simon 2010; Simon et al. 2013) to study a community consisting of a stochastically varying number *G* of multicellular groups. Within each group *i*, there are *N*_*i*_ individual cells, which may belong to any of *m* different species. Regardless of species, each cell is either wild-type (cooperator) or mutant (cheater). We denote the number of cells of species *j* ∈ (1, *m*) within group *i* as *N*_*j,i*_, so that 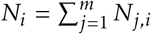. (Note that any particular group may contain fewer than *m* species; furthermore, species may go extinct, in which case the realized number of species in the community will be less than *m*.)

Community dynamics unfold simultaneously at both levels of biological organization: both the number of groups and the number of cells within each group vary over time. The number of cells within each group changes due to cell birth and cell death events, which occur at rates *b*_*j,i*_ and *d*_*i*_, respectively (section 2.1). Simulaneously, the groups themselves undergo fission (group birth) and extinction (group death) events, at rates *B*_*i*_ and *D*_*i*_, respectively (section 2.2). Finally, migration of cells between groups occurs at a rate *v*.

The individual-based simulation was implemented using a Gillespie algorithm (Gillespie 1976). The Python code (Van Rossum and Drake Jr 1995) used to create the simulations and the R code (R Core Team 2019) used to generate the figures are available at https://github.com/simonvanvliet/MLS_GroupDynamics.git.

### 2.1 Cell dynamics

#### Cell birth rate

A cell’s birth rate depends on cooperative interactions with other cells in the same group. At a cost *γ* to themselves, wild-type cells (cooperators) contribute to these interactions, increasing the birth rate of others in the group. However, mutations turn cooperators into cheaters, who reap the benefits of interactions without contributing (we ignore back-mutation). As a result, mutants always multiply faster than conspecific wild-type cells in the same group, but their presence slows the overall growth of the group.

We consider two possibilities: single-species systems (*m* = 1), where groups resemble multicellular organisms, and multispecies systems (*m >* 1), where groups are akin to mixed biofilms. When *m* = 1, cooperation occurs between individuals of the same species, whereas for *m >* 1, cooperation occurs across species, thus resembling obligate mutualistic interactions.

Let 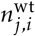 denote the number of wild-type cells of species *j* in group *i* (so that the number of mutants is 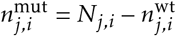. Then, the per capita realized birth rate of a mutant cell is equal to

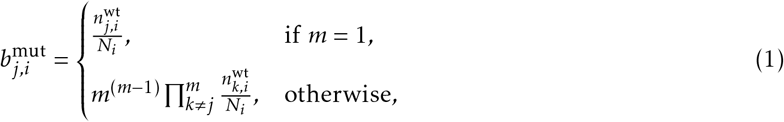

where the term *m*^(*m*−1)^ ensures that the total number of cells at equilibrium is the same across different values of *m*. Eq. 1 means that, when *m* = 1, a cell can only grow in the presence of conspecific cooperating partners, and when *m >* 1 a cell of species *j* can only grow in the presence of cooperating partners of all *m* − 1 other species (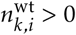 for all *k* ≠ *j*). Note that, when *m >* 1, growth does not depend on the presence of conspecifics, because it is meant to represent mutualistic interactions such as cross-feeding. The realized birth rate of a wild-type cell is then 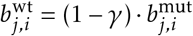, where *γ* is the cost of cooperation. Whenever a new cell is born, it stays in the same group as the parent. If a wild-type cell gives birth, with probability *μ* it generates a mutant cell of the same species and with probability (1 − *μ*) the offspring remains wild-type. We only consider mutations from wild-type to mutant in same species: there is no back-mutation or mutations across species.

#### Cell death rate

The per capita death rate *d*_*i*_ for all cells in group *i* is given by:

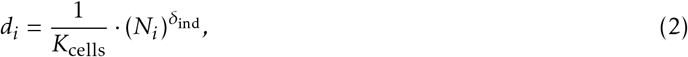

where *K*_cells_ is the within-group carrying capacity, and *δ*_ind_ is an indicator variable that can take the values 0 or 1. If *δ*_ind_ = 1, death rate increases linearly with group size, thus ensuring density regulation within the group; if *δ*_ind_ = 0, death rate is constant (no density dependence).

### 2.2 Group dynamics

#### Group fission rate

We use the term “fission rate” to refer to the rate at which groups reproduce (to avoid confusion with the birth of individual cells). All else being equal, groups with higher fission rates are favored by group selection. We consider the possibility that bigger groups are more likely to reproduce: the rate increases (with constant of proportionality *S*) with *N*_*i*_, up from a minimum value of *B*_0_. In summary,

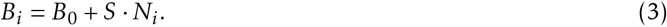

#### Modes of fragmentation

When a group of size *N*_*i*_ reproduces, it fissions into a parental group and one or more offspring groups, according to its mode of fragmentation. We assume that the offspring size is homogeneous, i.e., all offspring groups have the same expected size (this reduces the space of possible modes of fragmentation when compared to the more exhaustive approach used by Pichugin et al. 2017). Our model accommodates a wide variety of modes, without attempting to be exhaustive. To do this, we consider a triangular space of fragmentation strategies that is defined by two parameters (Fig. 1).

The first parameter (0 *< n* ≤ 1) determines the expected *number* of cells that are transmitted to offspring, as a fraction of the parent’s cell number. Whenever a group reproduces, we first draw this number of cells, *N*_offspring_, from a Poisson distribution with expectation *n* · *N*_*i*_, truncated at *N*_*i*_. The second parameter (0 *< s* ≤ 0.5) determines the expected *size* of each offspring group, again as a fraction of the parent’s cell number. This size, *σ*_offspring_, is drawn from a Poisson distribution with expectation *s* · *N*_*i*_, bounded between 1 (the size of a single cell) and *N*_offspring_. From these two quantities we then calculate the total number of offspring groups, *G*_offspring_ = ceil(*N*_offspring_/*σ*_offspring_). The first *G*_offspring_ − 1 offspring groups are all assigned *σ*_offspring_ cells, which are randomly sampled (without replacement) from the parent group. The final off-spring group is assigned the remaining cells, again by random sampling (without replacement) from the parent group. Finally, we remove all cells that are transmitted to the offspring groups from the parent group. The parameter *s* is analogous to the propagule size parameter in Kondrashov (1994) and Roze and Michod (2001). The only valid combinations of parameters obey the condition *s < n <* 1 − *s*. The lower triangle is excluded because it is logically impossible, whereas the upper triangle is excluded because one of the groups that result from the fissioning process is arbitrarily labelled as the parent group.

We refer to the three corners of this parameter space as *archetypal* fragmentation modes (Fig. 1): in *binary fission*, the parent group divides into two equally-sized groups; in *single-cell reproduction*, the parent group produces a single offspring group by releasing a unicellular propagule; finally, in *complete fragmentation*, the entire parent group disperses into its constituent cells. Every mode of fragmentation described in section 1 can be accommodated in this parameter space.

#### Group extinction rate

We use the term “extinction rate” to refer to the rate at which groups die (to avoid confusion with the death of individual cells). The extinction rate of group *i* can in general depend on the number of cells in the group *N*_*i*_, the number of cells in the entire community *N*_tot_ = Σ_*i*_*N*_*i*_ (which we also refer to as “community productivity”), the number of groups *G*, or some combination of these. The three different types of density-dependence are turned on or off by the indicator parameters *δ*_cells_, *δ*_tot_, and *δ*_groups_, respectively. This results in the per capita extinction rate

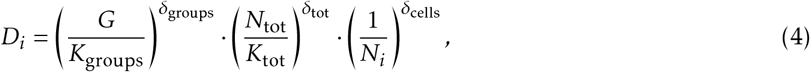

where the parameters *K*_groups_ and *K*_tot_ roughly scale the total number of groups and/or cells in the community. In the main manuscript, we will focus on the case in which *δ*_groups_ = 0, *δ*_tot_ = 1, and *δ*_cells_ = 0, which means that the extinction rate increases with the total number of cells in the community (*N*_tot_). We explore other choices of values for the indicator parameters in section S2. Groups also go extinct whenever the number of constituent cells drops to zero, e.g., due to stochastic sampling.

#### Migration rate

Each cell has a per capita migration rate *v*. Whenever a cell migration event occurs, the migrant cell leaves its group and joins a second, randomly chosen group.

### 2.3 Evolution of the fragmentation mode

To study the multilevel evolutionary dynamics of fragmentation mode, we characterize every cell in the population with a quantitative phenotype vector (*s, n*). These traits could represent, for example, the properties of the materials that are excreted into the extracellular matrix. The average trait value of the group determines the group’s position in state space (Fig. 1) and hence its mode of fragmentation. Thus, the properties of the individual cells give rise to an emergent group property. When cells reproduce, their offspring inherit their parent’s trait values; however, with a small probability *μ*_*s*_ or *μ*_*n*_, small-effect mutations may occur. For computational efficiency, we discretize the phenotype space and assume that mutations always occur between adjacent phenotype bins. We do not allow mutants outside the bounds of the phenotype space: any such mutants are moved along the *n*-direction and placed on the boundary of the allowed space.

## 3 Results

### 3.1 Multilevel selection can maintain cooperators and avert mutational meltdown

We will first consider groups consisting of a single species. When there are no mutations, the strategy of complete fragmentation maximizes equilibrium population size, measured either as total number of groups (Fig. 2A) or as total number of cells (i.e., productivity, Fig. 2B). But what happens in the presence of mutations?

**Figure 2:**
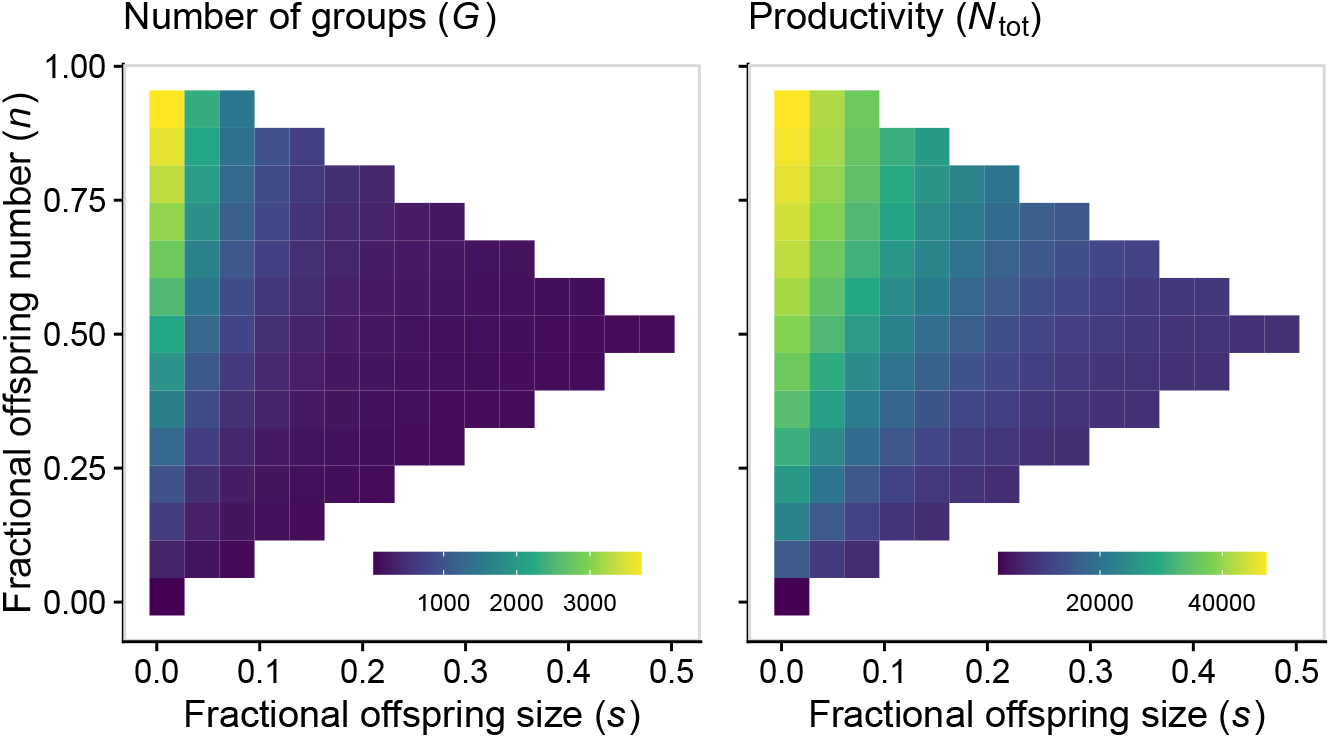
In the absence of mutations, with a single species, complete fragmentation into single cells maximizes equilibrium population size, measured either as total number of groups (*G*) or as total productivity (*N*_tot_). Parameters: *μ* = 0; all other parameters set to the default values (table S1).

Because mutations constantly create cheaters, we expect slower growing wild-type cells to go extinct over evolutionary time in the absence of group-level events. Furthermore, because all cells depend on the presence of cooperators to reproduce, this process is expected to lead to mutational meltdown: the extinction of the entire community due to the influx and spread of mutations. Indeed, when we set group-level rates to zero, we observe the accumulation of mutations, leading to the decrease in group size and eventual extinction of the community (Fig. 3A and black lines in Fig. 3B, C).

**Figure 3:**
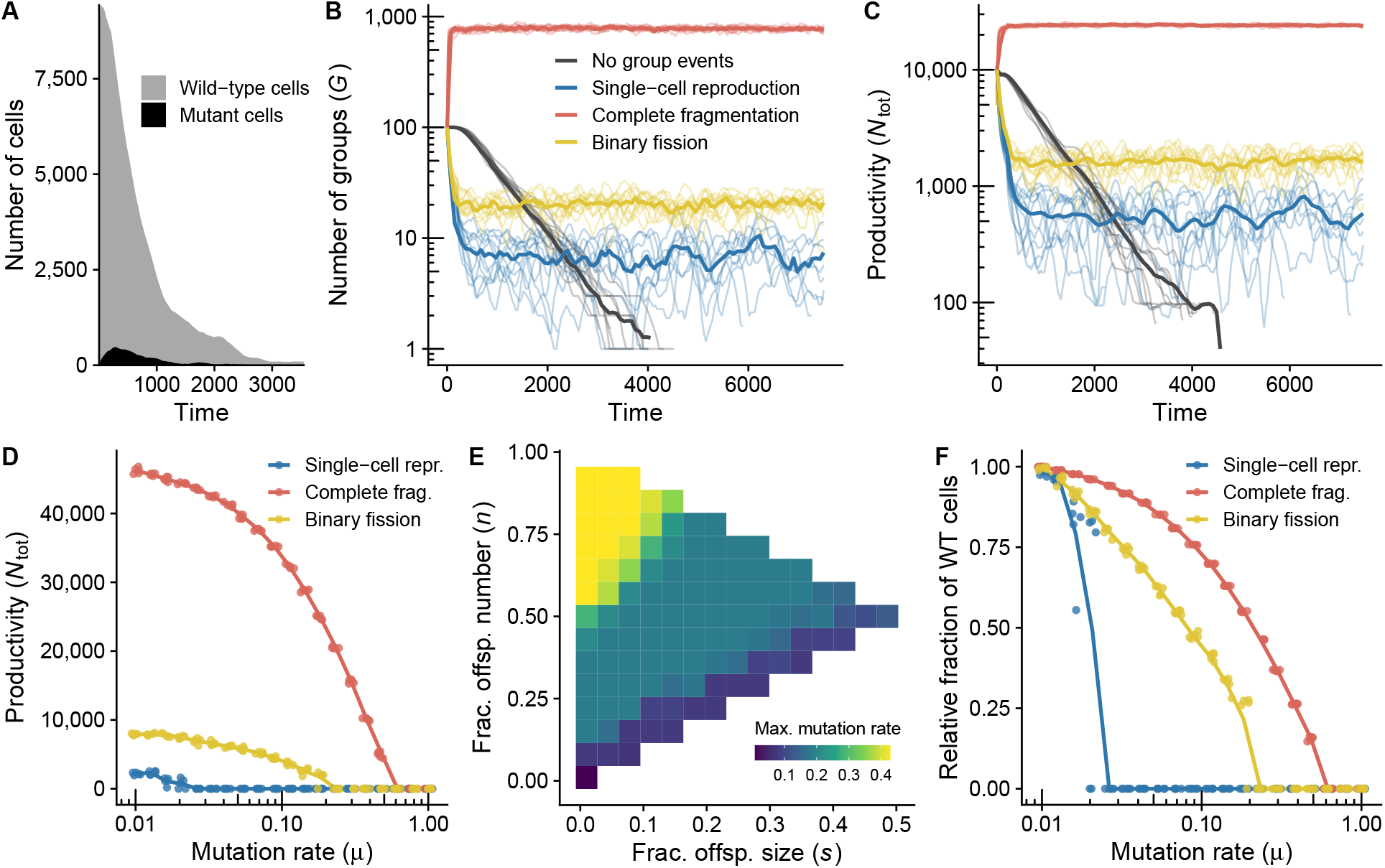
When there are no group events, mutations cause community extinction, but group-level events prevent this fate, as exemplified by the three archetypal modes of fragmentation. **A:** Example replicate with no group events, showing fast initial rise in the number of mutants and the consequential decrease in the total cell number, resulting in extinction. **B, C:** Temporal dynamics of the number of groups (**B**) and total productivity (**C**) for different reproduction modes. Thin lines are moving averages of individual replicates; solid lines are averages across replicates. **D:** By simulating population dynamics for various values of the mutation rate (*μ*), we can identify the value at which the population undergoes mutational meltdown–driven extinction (i.e., that strategies’ maximum mutation rate). **E:** For every point in the strategy space, we calculated the maximum mutation rate (shown in panel **D** for the three archetypes). **F:** The percentage of wild-type cells at equilibrium decreases with mutation rate (*μ*); the plot depicts the fraction of wild-type cells relative to the equilibrium fraction when *μ* is very small (*μ* = 0.01). Parameters: for the lines with group events, *B*_0_ = 0.01; all other parameters set to the default values (table S1).

However, group level selection can maintain cooperators and prevent meltdown. Cells in groups with many cooperators multiply faster than cells in groups with few cooperators. Furthermore, when these groups fission, they give birth to offspring groups that also have higher frequency of cooperators. Bigger groups are also less likely to collapse due to the stochastic death of all its member cells (driven by the accumulation of mutations and by genetic drift), and will therefore remain in the population for longer and have more opportunities to fission. Accordingly, when group-level rates are nonzero, selection at the level of the group can avert mutational meltdown (Fig. 3B, C).

### 3.2 Complete fragmentation minimizes mutation load

Previous theoretical work (Bergstrom and Pritchard 1998; Grosberg and Strathmann 1998; Kondrashov 1994; Roze and Michod 2001) predicts that, in single-species systems, small offspring sizes should be most resilient against mutational meltdown, since single-cell bottlenecks expose harmful mutations to natural selection. In agreement with these predictions, we found that the mode of group fragmentation has important consequences for the capacity of the population to persist in the presence of mutations (Fig. 3D). In single-species communities (*m* = 1), the complete fragmentation archetype and strategies close to it are able to avoid mutational meltdown-driven extinction even for relatively high rates of mutation when compared to other strategies (Fig. 3D, E). This is because, although the average frequency of wild-type cells per group always decreases with mutation rate, the relative decrease is smaller for this archetype (Fig. 3F), as would be expected given the role of tight bottlenecks in eliminating deleterious mutations.

### 3.3 Multispecies communities are more vulnerable to mutational meltdown

We have seen that, when single-species groups reproduce by complete fragmentation, they are most resistant to mutational meltdown. Multispecies communities, in contrast, cannot resort to complete fragmentation, since in small fragments there is a high chance that one or more heterospecific types are missing. In the extreme case of unicellular fragments, the offspring cell can never grow because of the absence of mutualistic interactions. In the absence of mutations, multispecies communities are thus most productive when groups have larger offspring, which allows offspring to maintain a variety of species. In other words, for increasing species number, the productivity maximum moves rightward along the upper diagonal of the phenotype space (Fig. 4A, B).

**Figure 4:**
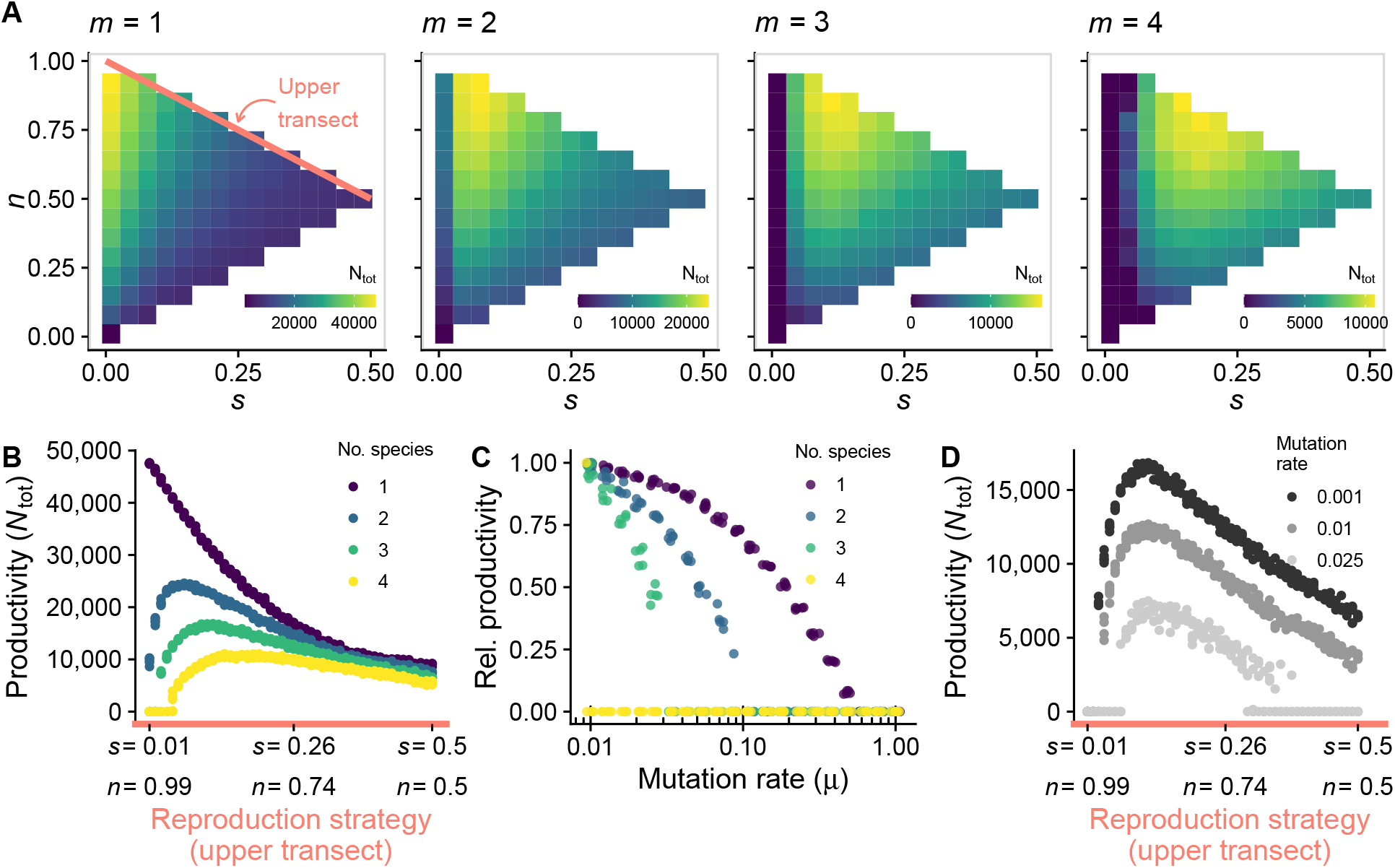
When community complexity is high, the productivity peak shifts away from unicellular bottlenecks. **A:** The color indicates equilibrium community productivity (*N*_tot_). As the number of species (*m*) increases, the strategy that maximizes *N*_tot_ moves rightward along the upper transect (pink line) of the strategy space. **B:** Equilibrium productivity as a function of strategies along the upper transect of the strategy space (corresponding to the pink line from panel **A**; ranges from complete fragmentation, on the left of the *x* axis, to binary fission, on the right), for different numbers of species. **C:** Equilibrium productivity decreases with mutation rate (*μ*); this decrease is faster for higher number of species (exemplified here for *s* = 0.1, *n* = 0.9.) **D:** Some strategies that do well with small *μ* (large offspring) are not viable when *μ* is large (shown here with *m* = 3). Parameters: all parameters are set to the default values, unless otherwise indicated (table S1).

Because multispecies communities need larger offspring, they are also more vulnerable to mutants (Fig. 4C). Therefore, there is a trade-off between resistance to mutational meltdown (low offspring size) and the maintenance of mutualistic interactions (high o spring size, Fig. 4D).

### 3.4 Size-dependent fragmentation rate and migration prevent mutational meltdown in multispecies communities

We have seen that (consistent with previous studies) multicellular organisms can reduce mutation load and prevent mutational meltdown by reducing propagule size (section 3.2), but that this strategy is not available in multispecies communities, since it deprives cells in daughter groups of their mutualistic partners (section 3.3). What reproductive strategies, then, allow for more complex multispecies communities to resolve this trade-off?

One alternative is size-dependent fragmentation rate. Larger groups may be more likely than smaller ones to undergo fission and thus produce offspring groups (in our model, this is achieved by increasing the slope parameter *S* in Eq. 3). Because all groups have equal extinction rate, groups with higher fission rate are favored by group selection. Hence, increased size-dependence in fissioning rate intensifies the effectiveness of group selection in purging mutants, since groups with many mutants grow slower and thus take longer to reproduce. By increasing the importance of group selection relative to individual-level selection, size-dependent fissioning should allow complex communities to withstand high rates of mutation. To test this hypothesis we calculated, for different values of *S*, the highest mutation rate that communities can withstand, over the entire strategy space. In other words: for each fragmentation mode we found the highest mutation rate that the population can handle before if collapses; we then selected the fragmentation mode for which this mutation rate is highest. We will later see that this strategy is also an evolutionary attractor, so we expect that this is the mode of fragmentation of groups at evolutionary equilibrium. The results confirm our prediction that at higher values of *S*, communities can survive in the presence of higher mutation rates (Fig. 5; see also Fig. S3 for a broader range of parameters).

**Figure 5:**
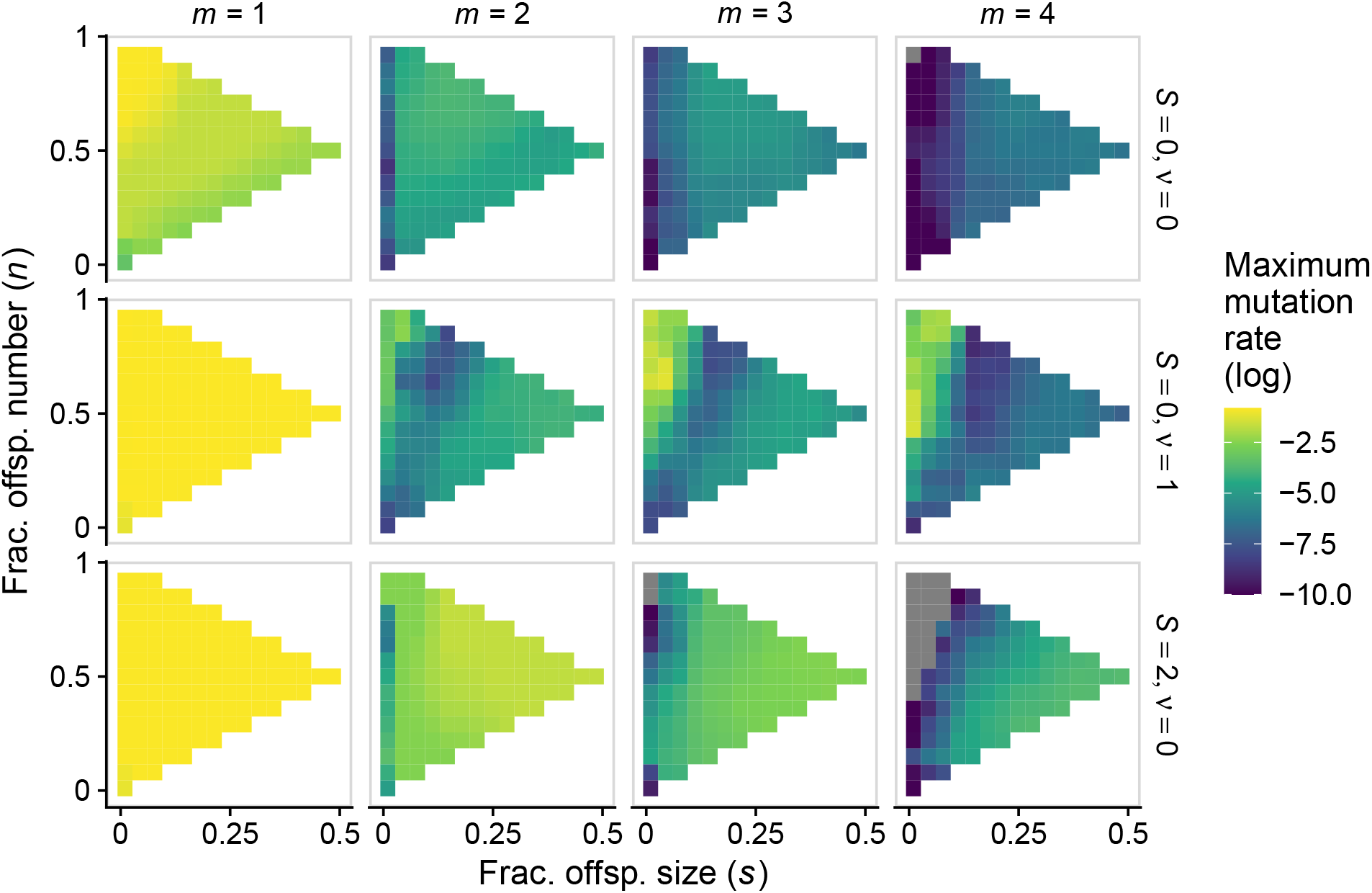
Migration and size-dependent fragmentation rate allow multispecies communities to resist mutational melt-down. Each panel shows, for each position in the strategy space, the maximum mutation rate a population can experience before going extinct (similar to Fig. 3E). Columns (from left to right) depict increasing number of species. Rows depict: no size-dependent fragmentation and no migration (*top*); migration and no size-dependent fragmentation (*center*); size-dependent fragmentation and no migration (*bottom*). Grey squares correspond to communities for which the maximum mutation rate is outside the range of our simulations or numerical errors. For each value of mutation rate, we assessed ten replicates per fragmentation mode; we then calculated the mean (across replicates) of the logarithm of maximum mutation rate. Each point in the figure corresponds to a nearest-neighbour average of this quantity. Parameters: all parameters are set to the default values, unless otherwise indicated (table S1). Figures S3 and S4 comprise a wider range of values of *S* and *v* and depict raw values of maximum mutation rate (rather than nearest-neighbour averaged values).

Another alternative is migration of cells between groups, which is common in bacterial biofilms. In multilevel selection theory, migration is often considered to be detrimental to group selection, because it decreases variance between groups and increases the role of individual-level selection. However, in multi-species systems, low to intermediate levels of migration can be beneficial, because they allow small offspring to recruit individuals from the environment and, thus, achieve the diversity necessary for mutualistic interactions (Fig. 5; see also Fig. S4 for a broader range of parameters).

Size-dependent fragmentation and migration are thus two possible solutions to the challenge of resisting the accumulation of mutations while also maintaining species diversity within groups. Size-dependent fragmentation increases the strength of group selection, which purges deleterious mutations, thus making small offspring sizes unnecessary. In contrast, migration allows for the recruitment of heterospecific cells, which allows diversity to be maintained even when offspring sizes are small.

### 3.5 Strategies that maximize community productivity are evolutionary attractors

Real communities vary in their mode of fragmentation. In fact, experiments in a variety of species— including *Chlamydomonas reinhardtii* (Herron et al. 2019; Ratcliff et al. 2013b), *Saccharomyces cerevisiae* (Ratcliff et al. 2013a), and *Pseudomonas fluorescens* (Hammerschmidt et al. 2014)—show that the mode of fragmentation can rapidly evolve under selection pressure.

Fragmentation mode has implications for quantities such as community productivity (total number of cells), number of groups, and total frequency of mutants in the community. However, different strategies may maximize each of these quantities. For example, when fragmentation rate is size-dependent, the number of groups is maximized when the slope of the fission function (*S*) is small (close to the complete fragmentation archetype), whereas the total community productivity is maximized when *S* is large (close to the binary fission archetype). Given that both the number of groups and the number of cells affect selection in different ways at both levels of organization, it is not obvious which mode of fragmentation is favored by natural selection under different conditions.

To answer this question, we simulated the multilevel evolutionary dynamics (section 2.3). Across a diverse set of parameters, we found that strategies that maximize community productivity (i.e., total number of cells) are evolutionary attractors and endpoints of evolution (for some examples, see Fig. 6). Therefore, when complexity is low, we expect evolution to lead to tight bottlenecks, but for more complex communities with more than one species, we expect group selection to promote the evolution of binary fission. When there are multiple species, the evolutionary outcome will depend on what mechanisms are in place to resolve the trade-off between maintaining complexity and eliminating deleterious mutations; for example, size-dependent fragmentation favours the evolution of binary fission (Fig. 6).

**Figure 6:**
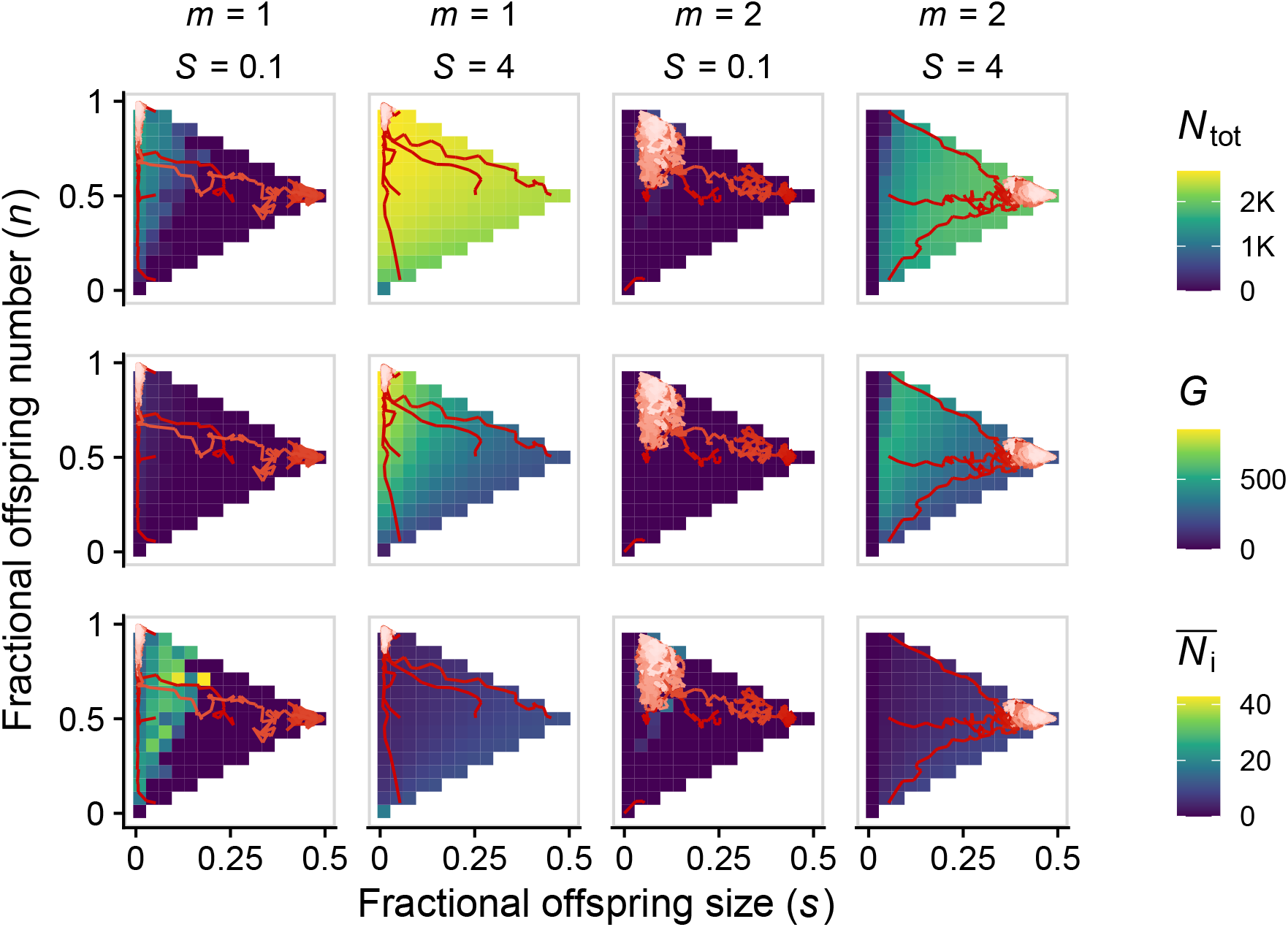
Evolution of fragmentation mode maximizes total number of cells (*N*_tot_, top row) rather than other quantities such as number of groups (*G*, middle row) or average group size (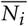, bottom row). Red lines represent evolutionary trajectories of the average mode of fragmentation over time (darker shades of red correspond to earlier time points). For each value of *S*, we assessed five initial phenotypes. Parameters: all parameters are set to the default values, unless otherwise indicated (table S1), except: *K*_cells_ = 100, *B*_0_ = 0.01, *K*_tot_ = 30,000 (when *S* = 0) or *K*_tot_ = 10,000 (otherwise), *μ*_*s*_ = *μ*_*n*_ = 10^−2^.

## 4 Discussion

Reproduction is the defining characteristic of life, yet organisms across the tree of life reproduce in many different ways. In organisms that have single-celled gametes, there is less within-organism variation than in organisms with many-celled propagules, which increases the strength of natural selection in removing deleterious mutations (Bergstrom and Pritchard 1998; Grosberg and Strathmann 1998; Kondrashov 1994; Roze and Michod 2001). Hence, single-celled bottlenecks allow populations to reduce mutation load. In this paper, we studied how this process affects the mode of fragmentation of multispecies collectives of organisms, such as microbial biofilms. We found that complex communities face a reproductive dilemma. On the one hand, their persistence relies on aligning the Darwinian interests of the group and its individual cells, which can be achieved by small reproductive bottlenecks (not unlike single-species organisms). On the other hand, due to stochastic sampling, small daughter groups lack the species diversity that multi-species communities rely on for ecological success. There is a tug-of-war between two competing selection pressures affecting daughter group size: maintaining species diversity while reducing mutation load. We explored two alternative solutions to this dilemma: migration (which makes small groups viable) and size-dependent fragmentation (which reduces load even within large groups).

Migration of cells between groups allows small groups to acquire species diversity. Due to stochastic sampling, some newly born groups are free of mutants, but may lack mutualistic partners. Thanks to migration, these groups will be able to recruit individuals of other species, whose presence is necessary for cell growth. Mutants also migrate, which may be detrimental for some groups. As long as the combined stochastic processes of birth and migration create some groups that have all species but no mutants, those groups will be favored by group-level selection. In nature, many mixed biofilms grow by a mechanism of “co-colonization” that resembles this process: one species often plays the role of the initial colonizer and other, mutualistic species later join (Elias and Banin 2012).

Strategies that intensify the effectiveness of group-level selection relative to individual-level selection can provide alternative ways to eliminate mutant cells. If larger groups are more likely than smaller ones to fission and produce offspring, group size becomes favored by multilevel selection. Since groups with fewer mutants grow faster, size-dependent fragmentation rates allow large, species-diverse groups to persist in the face of high mutation rates.

Migration and size-dependence are only two plausible solutions to the dilemma of fragmentation in multispecies communities. They are instructive in that each resolves the dilemma by relaxing one of the two conflicting selection pressures, but alternative solutions are, of course, entirely possible. One example is selective group extinction, which could have identical effects to selective group fission. Another example is the segregation of cooperators and defectors during fission events, also known as associative splitting (Bowles 2006; Eshelt and Cavalli-Sforza 1982; Haldane 1932; Hamilton 1975; Simon and Pilosov 2016; Wright 1943). In fact, when Kondrashov (1994) first proposed that small propagule size reduces mutation load, he pointed out that the effect will be most effective when mechanisms are in place to ensure that the propagules are as homogeneous as possible. Hence, mutation load is minimized when only very related cells are recruited to form a propagule (i.e., associative splitting). Multicellular collectives are able to increase relatedness between propagule cells by creating segregated germ lines. Such complex mechanisms, however, are not required to achieve associative splitting. Even simple aggregations of cells can ensure that their propagules are maximally related by maintaining spatial structure. If daughter cells remain close to their parents, then group fragments that break off from the mother group will be highly homogeneous. This process could potentially allow large daughter groups to eliminate mutation load, although such groups would presumably also be homogeneous in terms of their species composition. Another simple mechanism is differential adhesion, where each species remains tightly attached to one (or very few) of the other species, which could ensure multispecies propagules.

By incorporating migration and size-dependent fission rates, we observed that cooperative interactions can be maintained even for multi-species communities and for very high cooperation costs and mutation rates. Both migration and size-dependent fission are simple and realistic processes that are present in many natural microbial communities. Hence, our model suggests that cross-species cooperative interactions could potentially be stable in natural systems. This result stands in contrast to the view that cooperation is likely to be unimportant in microbial communities (e.g., Foster and Bell 2012).

The results we discussed are facilitated by the stochastic nature of our model. When newly born, small groups are created, sampling variation allows for the birth of groups that consist entirely of wild-type cells. In an infinite-size continuous limit, there would *always* be some fraction of mutants in every group. Because mutants grow faster than wild-type cells, even small fractions of mutants could pose challenges to the persistence of the community.

Our model also encompasses single-species groups, and we can contrast it to previous studies of fragmentation mode. We recover the classic result of Kondrashov (1994) (which has since been expanded by many other authors, e.g. Bergstrom and Pritchard 1998; Grosberg and Strathmann 1998; Roze and Michod 2001), viz. that by producing small propagules, single-species groups are able to persist in the presence of high mutation rates. We can also recover the result from Pichugin et al. (2019), Pichugin and Traulsen (2018), and Pichugin et al. (2017) who used an analytical model to calculate the optimal fragmentation mode for simple single-species groups in the absence of mutations. Specifically, we recover their main finding that the optimal fragmentation mode depends on the rate functions used for cell birth and death (Fig. S5). The main strength of our multilevel selection framework is that it is highly versatile: not only can we can combine these different strands of the previous literature in a single mathematical framework, but in addition we can use it to study multi-species groups, a topic which has received little attention so far. The versatility of our model also makes it easy to extend it in future work to further investigate how more complicated rate functions (e.g., nonlinear size-dependence in fission rate or different types of fission-associated costs) can explain the variety of fragmentation modes in nature.

## Acknowledgments

The authors are thankful to all members of the Doebeli lab for fruitful discussions. md, gjbh, and svv were funded by nserc Discovery grant no. 219930, awarded to md. svv was also funded by an Early Postdoc Mobility Fellowship from the Swiss National Foundation (grant no. 175123). gjbh received additional funding from ubc Faculty of Graduate and Postdoctoral Studies.

## Supplemental Information

### S1 List of parameters

Table S1 lists the parameters used in the model and their interpretation, and indicates the default values used for producing all figures (unless otherwise indicated in the text).

### S2 Other types of density regulation

As the community grows, the extinction rate (Eq. 4) increases, which maintains the number of cells bounded at some upper limit. In the main text, we focused on the case in which *δ*_tot_ = 1 and *δ*_groups_ = 0, meaning the the extinction rate increases with the number of *cells* in the community. Alternatively, it is conceivable that rate could instead increase with the number of *groups* in the community, in which case *δ*_tot_ = 0 and *δ*_groups_ = 1. This kind of density-dependence provides qualitatively similar results to the one explored in the main text. For example, it is still the case that increasing the number of species moves the productivity optimum away from complete fragmentation and toward binary fission, reflecting a trade-off between resistance to mutational meltdown at low offspring sizes and the maintenance of mutualistic interactions at high offspring sizes (Fig. S1).

**Table S1:** List of parameters and default values that were using for producing figures.

| Parameter | Default value |
| --- | --- |
| *Parameters affecting community complexity* |  |
| $m$ Initial (maximum) number of species | 1 |
| *Parameters affecting mutation and migration* |  |
| $\gamma$ Cost of cooperation | 0.01 |
| $\mu$ Rate of mutation | 0.001 |
| $\nu$ Rate of migration | 0 |
| *Parameters affecting within-group density regulation* |  |
| $K_{\text{cells}}$ Within-group carrying capacity | 100 |
| $\delta_{\text{ind}}$ Within-group density-dependence (indicator variable) | 1 |
| *Parameters affecting group fission rate (GFR)* |  |
| $B_0$ Minimum GFR | 0.05 |
| $S$ Slope of group size-dependence for GFR | 0 |
| *Parameters affecting group extinction rate (GER) and between-group density regulation* |  |
| $K_{\text{groups}}$ Scales the total number of groups in the community | 0 |
| $K_{\text{tot}}$ Scales the total number of cells in the community | $10^5$ |
| $\delta_{\text{groups}}$ Dependence of GER on number of groups (indicator variable) | 0 |
| $\delta_{\text{tot}}$ Dependence of GER on the community's number of cells (indicator variable) | 1 |
| $\delta_{\text{cells}}$ Dependence of GER on the group's number of cells (indicator variable) | 0 |
| *Parameters affecting mode of fragmentation* |  |
| $n$ Fractional offspring number | (0, 1] |
| $s$ Fractional offspring size | (0, 0.5] |

**Figure S1:**
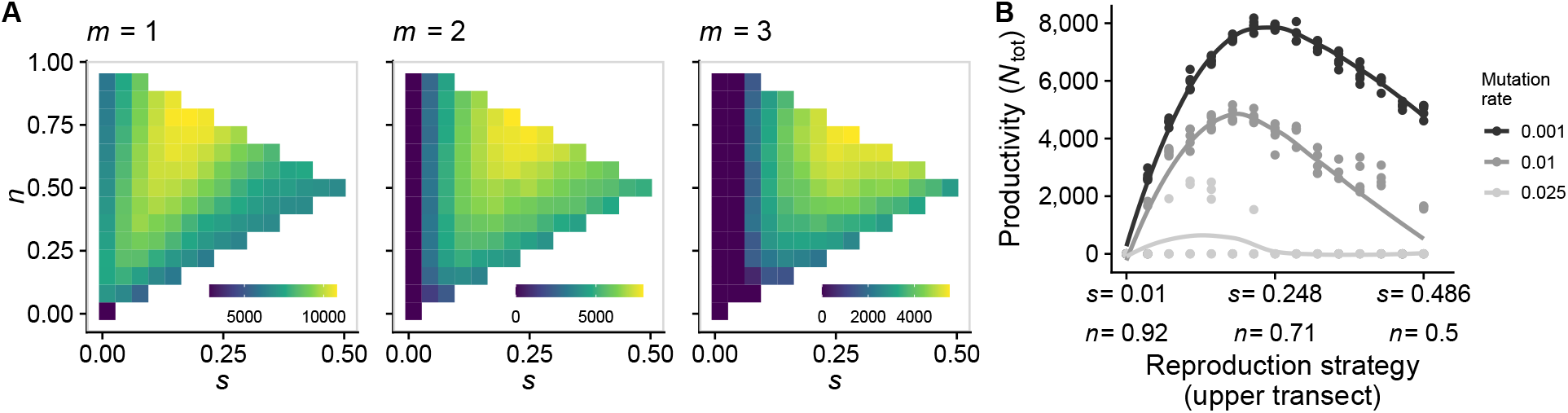
When extinction rate grows with number of groups instead of number of cells, the main qualitative model result does not change: increased community complexity shifts the productivity peak away from small bottlenecks and toward binary fission. **A:** The color indicates equilibrium community productivity (*N*_tot_). As the number of species (*m*) increases, the strategy that maximizes *N*_tot_ moves rightward along the upper transect of the strategy space. **B:** Some strategies that do well with small *μ* (large offspring) are not viable when *μ* is large (shown here with *m* = 2). Solid lines are LOESS smoothers. Parameters: *δ*_tot_ = 0, *δ*_groups_ = 1, *K*_groups_ = 103; all other parameters are set to the default values (table S1).

We also explored the possibility that the extinction rate increases not only with the number of cells in the community but, additionally, with the number of cells in the *group*, in which case *δ*_tot_ = 1, *δ*_cells_ = 1, and *δ*_groups_ = 0. In this case, larger groups are more likely to go extinct. In this case, the results are generally similar to the case in the main text; one difference is that, when complexity increases, the shift in the productivity peak away from small bottlenecks and toward binary fission is less pronounced. This makes sense, since there is a penalty on group size (Fig. S2).

### S3 Effect of migration and of size-dependent fragmentation on maximum mutation rate

In section 3.3, we showed that size-dependent fragmentation rate and migration can allow multispecies communities to prevent mutational meltdown (Fig. 5). The same results are shown in Fig. S3 for a broader range of values of *S* (slope of size-dependence), and in Fig. S4 for a broader range of values of *v* (migration rate).

**Figure S2:**
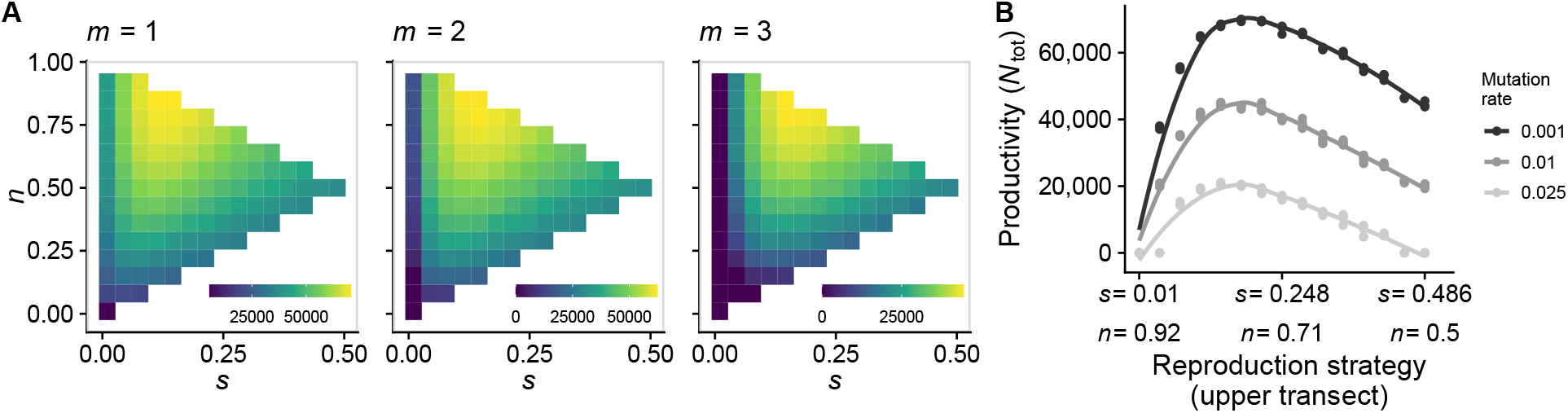
When extinction rate grows with group size in addition to number of cells, the effect of increased community complexity (i.e., the shift of the productivity peak away from small bottlenecks and toward binary fission) is less pronounced. **A**: The color indicates equilibrium community productivity (*N*_tot_). As the number of species (*m*) increases, the strategy that maximizes *N*_tot_ moves slightly rightward along the upper transect of the strategy space (shown here with *μ* = 0.01). **B**: Some strategies that do well with small *μ* (large offspring) are less viable when *μ* is large (shown here with *m* = 3). Solid lines are LOESS smoothers. Parameters: *δ*_cells_ = 1, *K*_tot_ = 10^4^; all other parameters are set to the default values (table S1).

### S4 Recovering the results of Pichugin et al. (2017)

In Pichugin et al. (2017), cells within group *i* grow with a birth rate *b*(*N*_*i*_) = 1 + *Mg*(*N*_*i*_), where *g*(*N*_*i*_) = [(*N*_*i*_ − 1)/(*N*_max_ − 2)]^*κ*^. Here, *N*_max_ is the maximum cell number in a group, *M* is the maximum benefit of group life, and *κ* is a “complementarity” parameter that measures how each additional cell increases the benefit. When the number of cells reaches *N*_max_, the group fissions according to a given fissioning strategy, or “partition”. Pichugin et al. (2017) tested all mathematically possible partitions for a given *N*_max_, and measured their group-level fitness. They found that the complementarity parameter *κ* is the main determinant of group-level fitness. Overall, if there are diminishing returns to the benefit of additional cells (*κ* ≪ 1), binary fragmentation is the most fit partition, but if there are increasing returns (*κ* ≫ 1), unicellular propagule production is the most fit. Other parameters, such as maximum benefit *M*, have a smaller influence on group-level fitness.

Our framework is general enough to accommodate Pichugin et al. (2017) as a special case. We set the group extinction rate to zero, and the birth rate of a cell in group *i* to *b*_*i*_(*N*_*i*_) = 1 + *g*(*N*_*i*_), with *g*(*N*_*i*_) = [(*N*_*i*_ − 1)/(*K*_ind_ − 2)]^*κ*^. Finally, we set the group fission rate to zero if *N*_*i*_ < *K*_ind_, else *B*_*i*_ = *B*_0_ = 10^6^. When we measure group growth rate we observe the same result as Pichugin et al. (2017) (Fig. S5A). The same pattern is observed when measuring cell growth rate (Fig. S5B), which makes sense because, as soon as group size has equilibrated, both cells and groups have to grow at the same rate.

**Figure S3:**
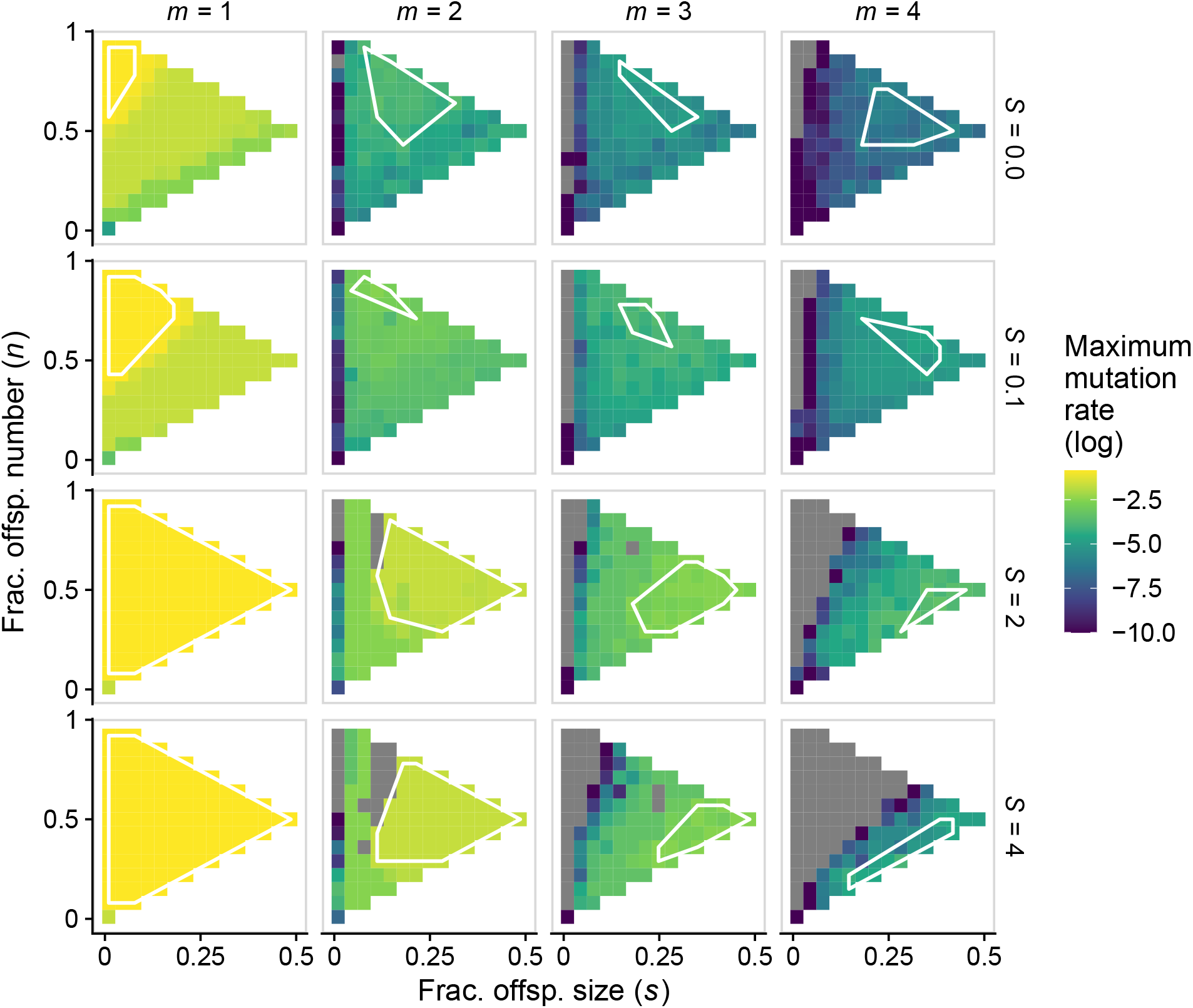
Size-dependent fragmentation rate allows multispecies communities to resist mutational meltdown. Each panel shows, for each position in the strategy space, the maximum mutation a population can experience before going extinct (similar to Fig. 3E and Fig. 5). Columns (from left to right) depict increasing number of species. Rows (from top to bottom) depict increasing slope of size-dependence. Grey squares correspond to communities for which the maximum mutation rate is outside the range of our simulations or numerical errors. The white polygons are convex hulls containing the five highest values of maximum mutation rate (to help guide the eye). For each value of mutation rate, we assessed ten replicates per fragmentation mode; the figure shows the mean (across replicates) of the logarithm of maximum mutation rate. Parameters: all parameters are set to the default values, unless otherwise indicated (table S1).

**Figure S4:**
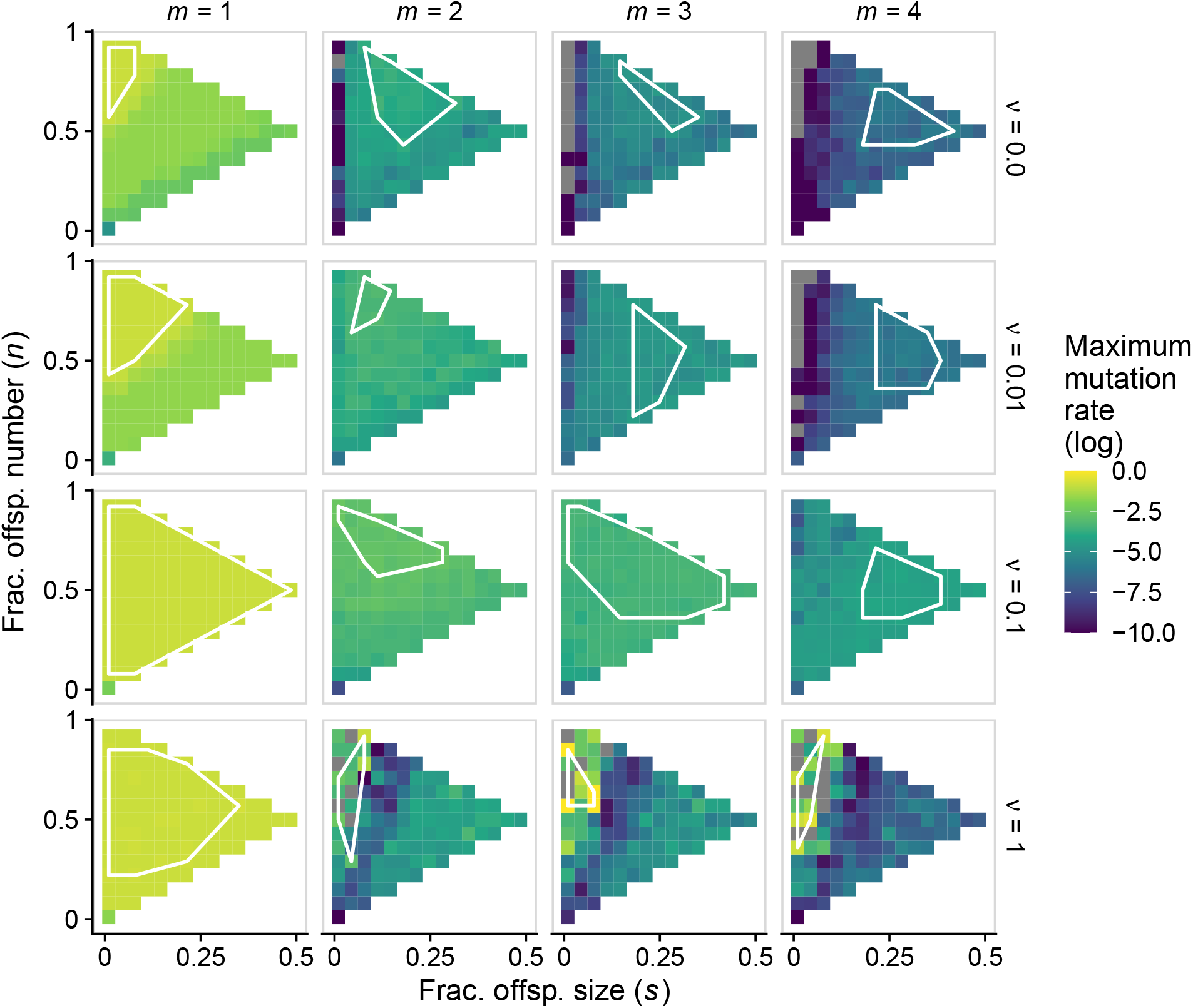
Low to intermediate migration rates allow multispecies communities to resist mutational meltdown. Each panel shows, for each position in the strategy space, the maximum mutation a population can experience before going extinct (similar to Fig. 3E and Fig. 5). Columns (from left to right) depict increasing number of species. Rows (from top to bottom) depict increasing migration rates. Grey squares correspond to communities for which the maximum mutation rate is outside the range of our simulations or numerical errors. The white polygons are convex hulls containing the five highest values of maximum mutation rate (to help guide the eye). For each value of mutation rate, we assessed ten replicates per fragmentation mode; the figure shows the mean (across replicates) of the logarithm of maximum mutation rate. Parameters: all parameters are set to the default values, unless otherwise indicated (table S1).

**Figure S5:**
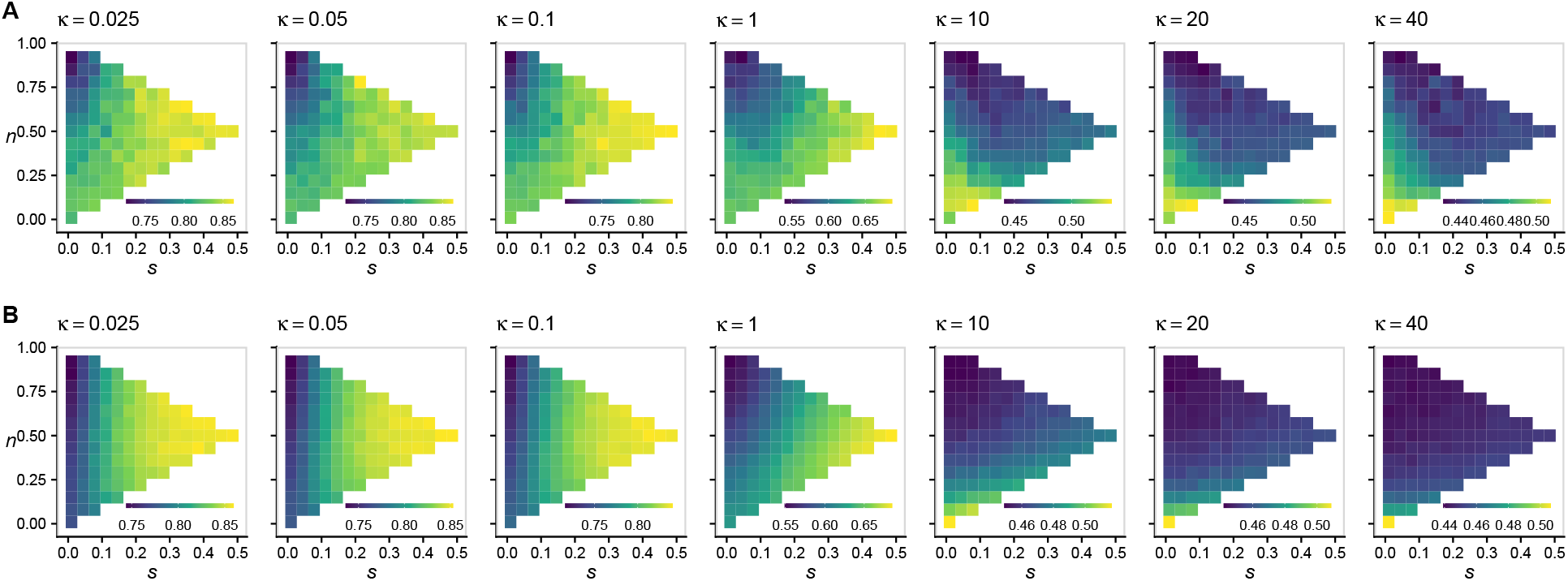
In the absence of group-level density-dependence, if the cell birth rate increases with group size, the complementarity parameter *κ* determines which strategy maximizes fitness—measured either as group-level growth rate (**A**) or as cell-level growth rate (**B**). For *κ* ≪ 1 (diminishing returns), binary fragmentation maximizes fitness, whereas for *κ* ≫ 1 (increasing returns), single-cell reproduction maximizes fitness. The color scale indicates growth rate.

